# Reference-Aided Full-length Transcript Assembly, cDNA Cloning, and Molecular Characterization of *Coronatine-insensitive 1b* (*COI1b*) Gene in Coconut (*Cocos nucifera* L.)

**DOI:** 10.1101/2020.11.23.395202

**Authors:** Frenzee Kroeizha L. Pammit, Anand Noel C. Manohar, Darlon V. Lantican, Roanne R. Gardoce, Hayde F. Galvez

## Abstract

In the Philippines, 26% of the total agricultural land is devoted to coconut production making coconut one of the most valuable industrial crop in the country. However, the country’s multimillion-dollar coconut industry is threatened by the outbreak of coconut-scale insect (CSI) and other re-emerging insect pests promoting national research institutes to work jointly on developing new tolerant coconut varieties. Here, we report the cloning and characterization of *coronatine-insensitive 1* (*COI1*) gene, one of the candidate insect defense genes, using ‘Catigan Green Dwarf’ (CATD) genome sequence assembly as reference. Two (2) splicing variants were identified and annotated – CnCOI1b-1 and CnCOI1b-2. The full-length cDNA of CnCOI1b-1 was 7,919 bp with an ORF of 1,176 bp encoding for a deduced protein of 391 amino acids while CnCOI1b-2 has 2,360 bp full-length cDNA with an ORF of 1,743 bp encoding a deduced protein of 580 amino acids. The 3D structural model for the two (2) isoforms were generated through homology modelling. Functional analysis revealed that both isoforms are involved in various physiological and developmental plant processes including defense response of plants to insects and pathogens. Phylogenetic analysis confirms high degree of COI1 protein conservation during evolution, especially among monocot species.

**Key Message:** This paper reports the molecular cloning and characterization of *corononatine-insensitive I* (*COI1)* gene in coconut using reference-guided transcript assembly approach. As a well-known insect defense-response gene in other crops, the results of this study are expected to assist in the development of new resistant coconut varieties as one of the strategies to address threats in coconut production.

## Introduction

More than 90 countries produce over 60M metric tons of coconut annually (FAOSTAT 2014). Currently, Philippines is the second largest producer of coconut around the world but remained as the top global producer of coconut oil (PSA 2017). However, the outbreak of coconut-scale insect (CSI) in 2014 which affected more than 2.6 million nut-bearing coconut trees threatened the coconut production in the country (PCA 2017). Various insect pest control methods were utilized in order to address the outbreak including physical, chemical, and biological control approaches.

Recombinant DNA technologies allow the effective cloning and characterization of candidate insect defense genes. However, most of the genes involved for insect defense in coconut have not been fully sequenced and characterized. One of the candidate insect defense response genes in coconut is *coronatine-insensitive 1* (*COI1*) gene which is reported to be involved in the jasmonate (JA) signaling pathway (Howe 2008). Jasmonate is known as an immunity hormone that promotes plant growth- and defense-related processes, such as, wounding and necrophitic pathogens (Zhai et al. 2017). In the JA signaling pathway, COI1 protein forms a complex with SCF (Skp/Cullin/F-box) to form an E3 ubiquitin ligase known as SCFCOI1 complex (Xu et al. 2002). This protein complex is necessary for the ubiquitination and subsequent degradation of JAZ proteins, transcription repressors in JA signaling pathway, causing the de-repression of transcription factors and activation of JA response genes. To date, the only identified jasmonate receptor is the COI1 protein (Yan et al. 2013). In *Arabidopsis*, *COI1* mutants were found to be insensitive to JA resulting in defects in pest and pathogen resistance (Feys et al. 1994; Mewis et al. 2005). COI1 plays a vital role in wound- and JA-signaling since it was found to be required for the activation and repression of genes associated with wounding and JA signaling (Devoto et al. 2004). Similarly, COI1 in rice was reported to be indispensable in signaling component of rice defense response to chewing insects (Ye et al. 2012).

Several studies on molecular characterization of COI1 from dicots have been done but not much on monocots. In *Arabidopsis*, COI1 encodes 66 kDa protein (592 amino acid) containing N-terminal F-box motif and 16 LRRs which adopts the horseshoe-like shape (Xie et al. 1998; Katsir et al. 2008; Yan et al. 2013). There were three (3) closely related *COI1* homologs in rice (Lee et al. 2013) while only a single copy was identified in rubber tree (Peng et al. 2009). COI1 protein in *Aquilaris sinensis* is composed of 29.95% α-helix, 12.25% β-sheets, and 58.50% random coil (Liao et al. 2015).

With the advent of next-generation sequencing (NGS) and bioinformatics tools, the first whole genome sequence assembly of Catigan Green Dwarf” (CATD) variety of coconut has been generated using PacBio SMRT sequencing platform and 50X Illumina Miseq, and improved through Dovetail Chicago^®^ sequencing (Lantican et al. 2019). With the aid of the genome assembly and information from transcriptome data, full-length complementary deoxyribonucleic acid (cDNA) can be obtained. Furthermore, defense and host-response genes such as *COI1* can be characterized at the genome level with the aid of the genome assembly.

The main objective of this study is to clone and elucidate the molecular characteristics of *COI1* gene in coconut. Specifically, this study aims to (1) generate the full-length cDNA sequence of *COI1b* from transcriptome data in reference to ‘Catigan Green Dwarf’ (CATD) genome sequence assembly, (2) clone partial *COI1b* cDNA sequence to validate the predicted full-length cDNA sequence of *COI1b*, and (3) to characterize *COI1b* gene and obtain phylogenetic relationships of coconut COI1b against known COI1 protein sequences.

## Materials and Methods

### Generation of COI1b full-length cDNA sequences

The full-length assembly of *COI1b* transcript was generated using genome-guided transcript assembly approach. The gene sequence of *COI1* in *Elaeis guineensis* (NW_011551140.1) was downloaded from the database and used as input for BLASTn (Altschul et al. 1997) search for gene homolog in the existing build of coconut ‘Catigan Green Dwarf’ genome (Lantican et al. 2019). The *COI1*-harboring genomic scaffold of coconut was extracted using a local perl script (https://github.com/solgenomics/sgn-biotools/blob/master/bin/fasta_extract.pl) and utilized as reference in the genome-guided assembly of *COI1b* full-length transcript. Sequence Read Archive (SRA) files from ‘Chowghat Green Dwarf’ (CGD; SRR1173229), ‘Taro Green Dwarf’ (TGD; SRR1273070), ‘Aromatic Green Dwarf’ (AGD; SRR1273070), and ‘West Coast Tall’ (WCT; SRR1137438) coconut varieties were retrieved from NCBI and preprocessed using Trimmomatic v0.36 (Bolger et al. 2014; SLIDING WINDOW: 5:30 LEADING: 5 TRAILING:5 MINLEN:85) to remove low-quality base calls. The preprocessed RNA-seq reads were aligned to the *COI1b*-containing coconut genomic scaffold using TopHat2 (Kim et al. 2013) aligner with default parameters. The resulting binary map alignment (BAM) files from each data set were used as input to the genome-guided Trinity software module (Grabherr et al. 2011) to assemble the full length *COI1b* transcript expressed across the four (4) coconut varieties.

### Plant Material

Healthy and coconut scale insect (CSI)-infested coconut leaf samples were collected from CATD variety of coconut at the Institute of Plant Breeding, University of the Philippines Los Baños. Sterile distilled water and RNAse AWAY^TM^ (Invitrogen Corporation, USA) was used to wash sample surface and remove possible RNA-degrading contaminants.

### Extraction of total RNA

Approximately 250 mg of leaf tissue was used for total RNA extraction using PureLink^TM^ Plant RNA reagent following the manufacturer’s protocol (Invitrogen Corporation, USA). A total of 20 μL diethyl pyrocarbonate (DEPC) water was used to resuspend each sample. In order to ensure the purity of the samples, the isolated total RNA was treated with DNase (Invitrogen Corporation, USA) followed by 1% agarose gel electrophoresis in 1x TBE buffer (90mM tris-borate, 2nM EDTA) to check the RNA integrity. The quantity of the RNA was further determined using Qubit 2.0 Fluorometer (Life Technologies Corporation, USA). The first-strand cDNA was synthesized using Superscript III First-Strand Synthesis System (Invitrogen Corporation, USA) following the kit’s protocol.

### PCR Amplification

The mRNA sequence of *COI1* among different palm species was mined in the NCBI public repository (https://ncbi.nlm.nih.gov) and *Elaeis guineensis* coronatine-insensitve 1 homolog b (XM_010909322.2) cDNA sequence was retrieved. With the retrieved cDNA sequence, the gene-specific primers (GSPs) were designed using Primer3Plus software (Untergasser et al. 2012). The designed GSPs were subjected to *in silico* PCR using the CATD genome sequence assembly to determine complementation and specificity prior to primer synthesis. The primer pair exhibiting *in silico* amplification product with specificity to the target gene of interest was sent for outsourced primer synthesis (Invitrogen Corporation, USA).

The thermal cycler profile to amplify the mid-transcript of the target gene is as follows: 94°C for 2 min; 35 cycles of 94°C for 15 s, 62°C for 30 s, and 68°C for 1 min. The PCR reaction cocktail has a total reaction volume of 25 μL composed of 10X HiFi Buffer (1X), MgSO4 (2mM), dNTPs (0.2 mM), 0.2μM each of forward (5’-**GCATTGGAAGAGTTTGGTGGGGGCTCA**-3’) and reverse primer (5’-**TCCAGCTTCTGCAGGCTTGGGCAAC**-3’), Platinum^®^ *Taq* DNA Polymerase HiFi (1U), and cDNA (30ng). The PCR products were subjected to 1% agarose gel electrophoresis in 1x TBE buffer (90mM tris-borate, 2nM EDTA) for 40 min at 100V to validate fragment size, and specificity and quality of the amplification product. Once validated, amplified cDNA fragments were excised from the agarose gel and further purified using the PureLink^®^ Quick Gel Extraction Kit (Life Technologies Corporation, USA). The gel extracted cDNA fragments were eluted with DEPC water to a final volume of 35 μL.

### Gene Cloning

The mid-transcript PCR product was ligated to pGEM^®^-T Easy vector (Promega Corporation, USA), following the manufacturer’s instructions and transformed into *E. coli* JM109 competent cells. The transformants were screened through blue-white screening and T7/SP6 colony PCR. The positive transformants were grown overnight in Lauria-Bertani (LB) broth containing ampicillin (100μg/mL). The plasmids were isolated using the PureLink^®^ Quick Plasmid Miniprep (Life Technologies Corporation, USA) following manufacturer’s instructions. Then, 3 μL of the purified plasmid DNA recovered was subjected to plasmid PCR using T7/SP6 sequencing primers to a total volume of 10 μL in order to detect the integrity of the recovered DNA plasmids. High quality plasmid extracts were sent for double-pass outsourced capillary sequencing (AIT Biotech, Singapore).

Raw sequence files (.abl and .seq file) that were generated from the capillary sequencing were pre-processed based on the sequence alignment of the designed primer pairs using the Vector NTI software (Invitrogen, USA). The trimmed sequences were then loaded to the CAP3 Sequence Assembly Program (Huang and Madan 1999) in order to generate the contig assembly from the double-pass paired reads.

### Characterization of reference-aided full-length COI1b cDNA sequences and deduced amino acid sequences

Multiple sequence alignment (MSA) was performed on the assembled *COI1b* full-length assembled variants from CGD, TGD, AGD, and WCT and the corresponding CATD gene sequence of *COI1b* gene in CATD using Clustal Omega (Sievers et al. 2011). The structural gene annotations were manually done using SnapGene software (GSL Biotech; www.snapgene.com). Similarly, the cloned mid-transcript sequence was also aligned to the consensus sequences to further validate the *COI1b* gene identity. The cDNA sequences of *CnCOI1* were also compared with sequences found in the database using BLASTn (Altschul et al. 1997).

The deduced amino acid sequences of coconut COI1b were derived by translating the mRNA sequences at different reading frame using the ORF Finder (https://www.ncbi.nlm.nih.gov/orffinder/) and conserved domains were identified using NCBI Conserved Domain Database (https://www.ncbi.nlm.nih.gov/Structure/cdd/wrpsb.cgi).. The composition of the amino acid sequences, molecular weight, and isoelectric point (pI) were predicted and analyzed using ProtParam (https://web.expasy.org/protparam/). The 3D molecular models were constructed through homology modelling using I-TASSER (Zhang 2008; Roy et al. 2010; Yang et al. 2015) and functional analysis using COFACTOR and COACH (https://zhanglab.ccmb.med.umich.edu). The coconut COI1b amino acid sequences were also used for BLASTn search (Altschul et al. 1997) and retrieval of homologous COI1 proteins from other species in the NCBI database (https://ncbi.nlm.nih.gov). Multiple sequence alignment using Clustal Omega (Sievers et al. 2011) was also performed followed by phylogenetic tree construction using FastTree (Price et al. 2010) to establish protein divergence and relationships among the COI1 amino acid sequences.

## Results

### Generation and validation of full-length CO11b cDNA

Multiple sequence alignment of the *COI1b* full-length assembled transcripts showed two (2) splicing variant detected in CGD and TGD while only one (1) splicing variant for WCT and AGD. CGD assembled transcript 1 and TGD assembled transcript 2 have 7590 bp and 7919 bp transcript length, respectively while all the remaining transcripts are around 2000bp in length. All the varieties have 5’-3’ orientation except for WCT. Manual annotation from the transcript assembly showed that there are two (2) *COI1b* splicing variants in coconut (**Fig 1**).

**Fig 1.**
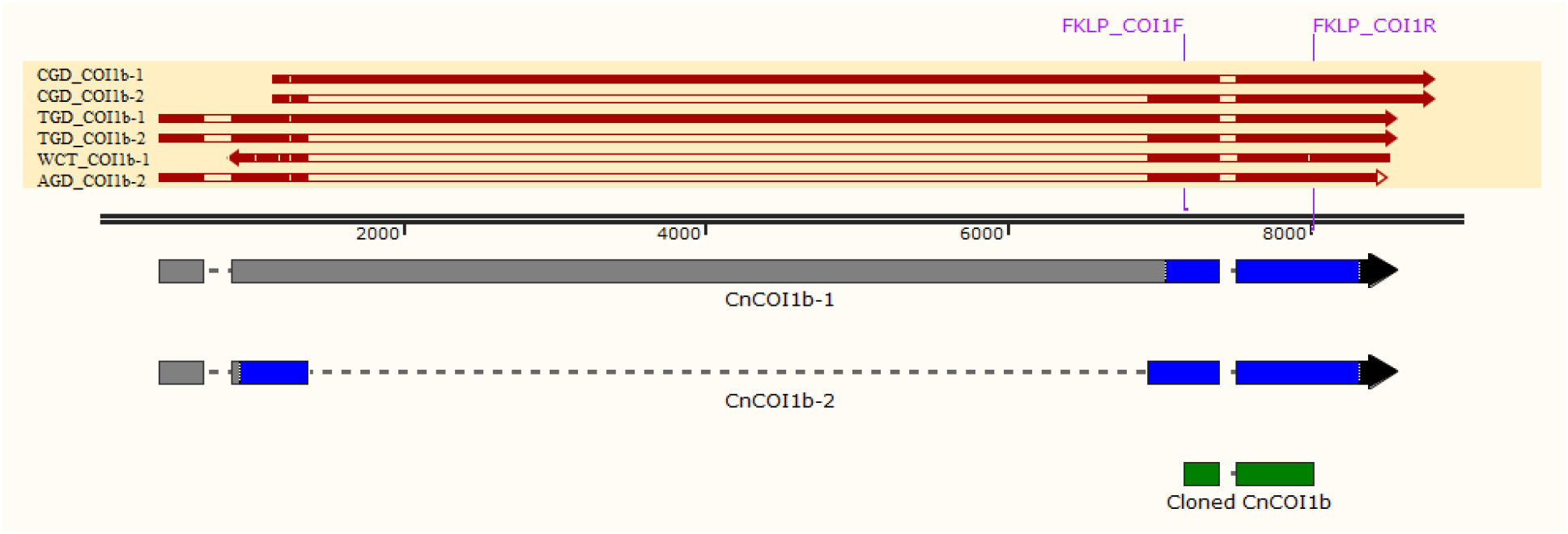
Visualization of the consensus cDNA sequences of CnCOI1b-1 and CnCOI1b-2 in reference transcript assembly. The arrow indicates the direction of transcription with the gene units indicated: 5’UTR (gray), ORF (blue), and 3’UTR (black).

The cloned mid-transcript sequence (752 bp; **Appendix 1**) of coconut *COI1b* gene was aligned to the two (2) *CnCOI1b* splicing variant in coconut generated using genome-guided transcript assembly approach (**Fig 1**). It covers approximately 10% of *CnCOI1b-1* cDNA (7,919 bp) and 32% of *CnCOIb-2* cDNA (2,360 bp). The cloned mid-transcript sequence also showed 96% similarity with *E. guineensis COI1b* as revealed by Blastn results. No sequence variation (**Appendix 2**) on the cloned mid-transcript was detected for healthy and insect-infested leaf samples.

### Characterization of COIb1 cDNA sequences

The full-length cDNA of the CnCOI1b-1 was 7,919 bp, containing a 1,176 bp ORF, with a 5’ UTR of 6,477 bp upstream the start codon and a 3’UTR of 266 bp downstream the stop codon (**Table 1**). On the other hand, CnCOI1b-2 has 2,360 bp full-length cDNA, containing a 1,743 bp ORF, with a 5’ UTR of 351 bp upstream the start codon and a 3’UTR of 266 bp downstream the stop codon. Both COI1b transcript variants in coconut have more than 8000 bp gene length. However, CnCOI1b-1 has significantly longer full-length cDNA, but shorter ORF compared to CnCOI1b-2. This observation is likely caused by the non-splicing of the intron 2 in the transcript variant CnCOI1b-1 relative to the sequence of the CnCOI1b-2. Upon further investigation within the sequence of the intron 2, several stop codons were found in the nucleotide stretch of the expressed sequence. Thus, it is classified as the 5’UTR of the longest ORF predicted by ORF Finder. Moreover, CnCOIb-2 and EgCOI1b have a very long 5561 bp intron which is absent in CnCOI1b-1, AtCOI, AsCOI1, and HbCOI1. Regardless, the cDNA of coconut COI1b sequences share 80.63% identical nucleotides (**Fig 2**) and more than 95% identical to the COI1b of cDNA sequence of *E. guineensis*.

**Table 1.**
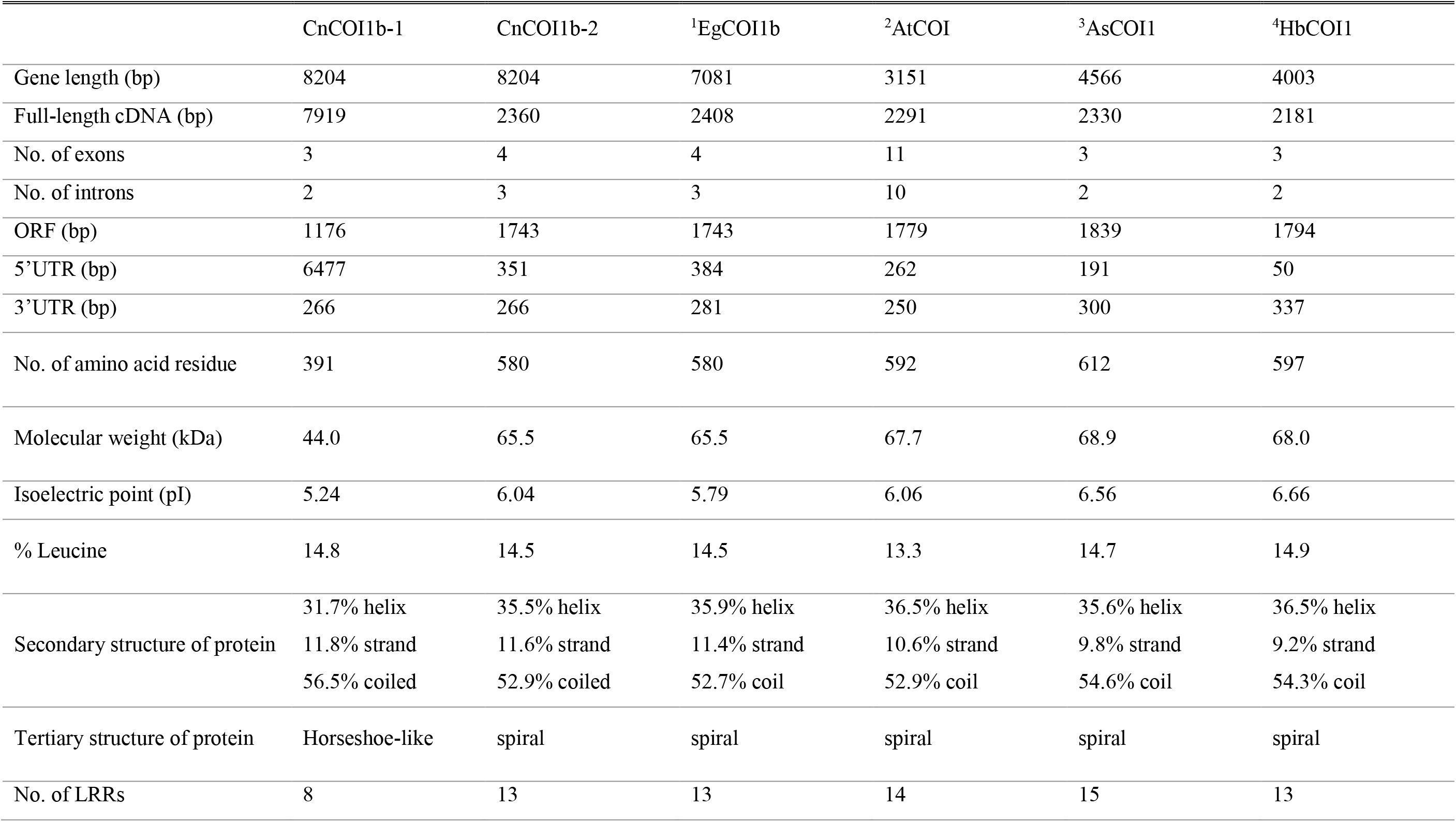
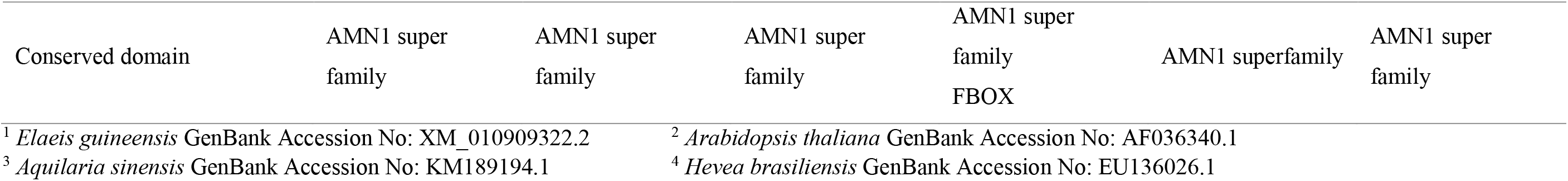
Comparison of cDNA and deduced amino acid sequences among different plant species.

**Fig 2.**
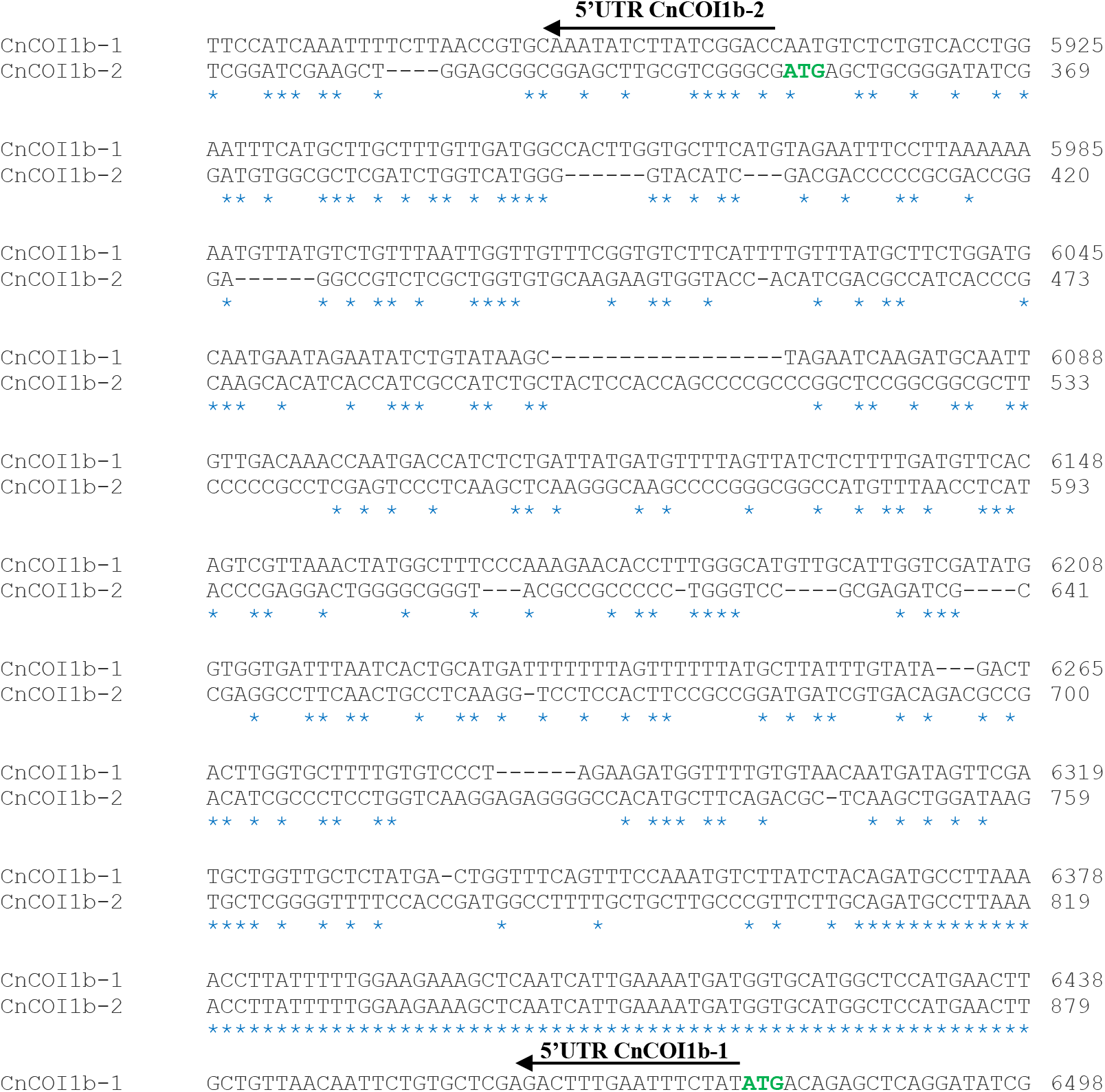
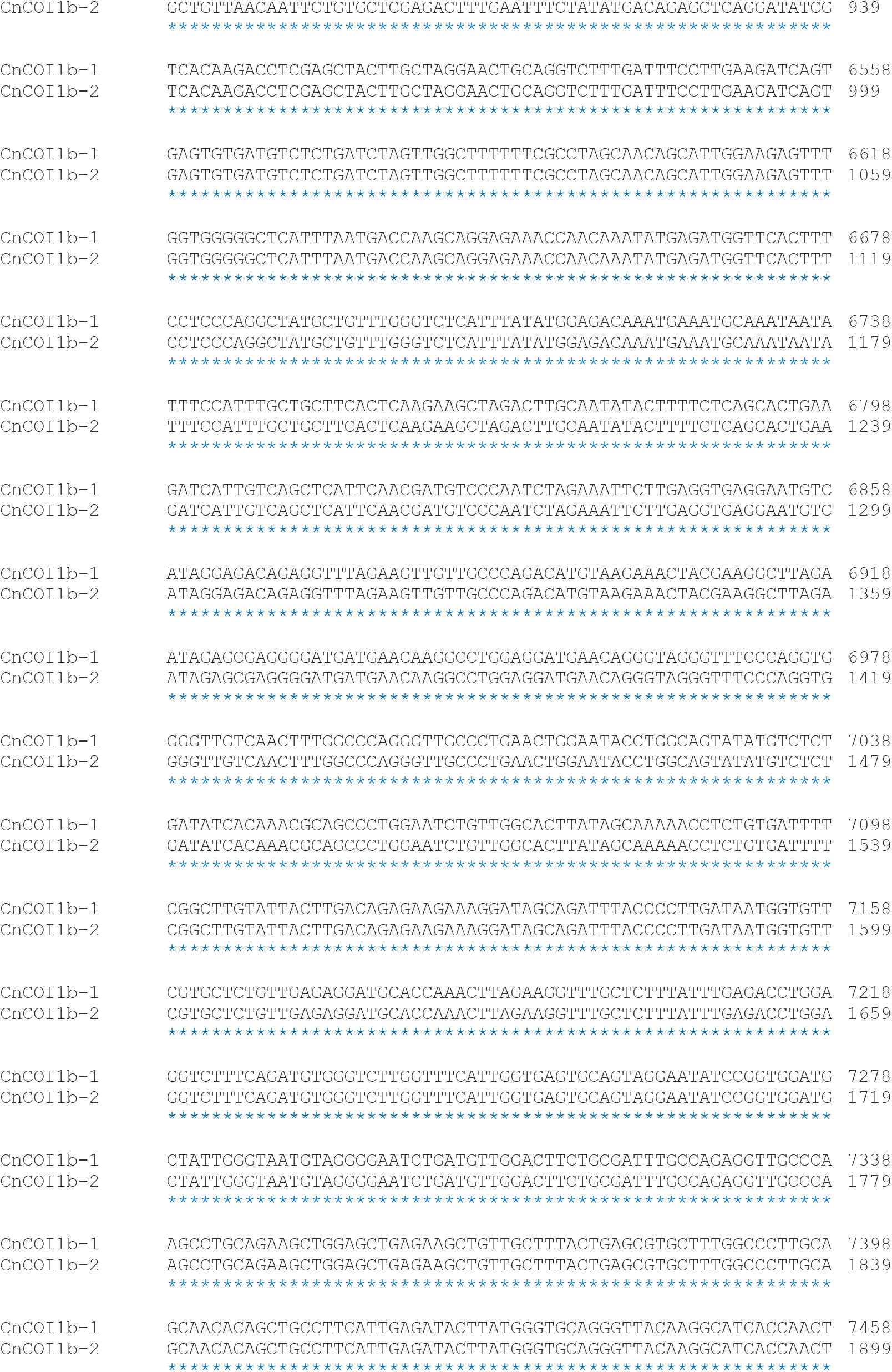
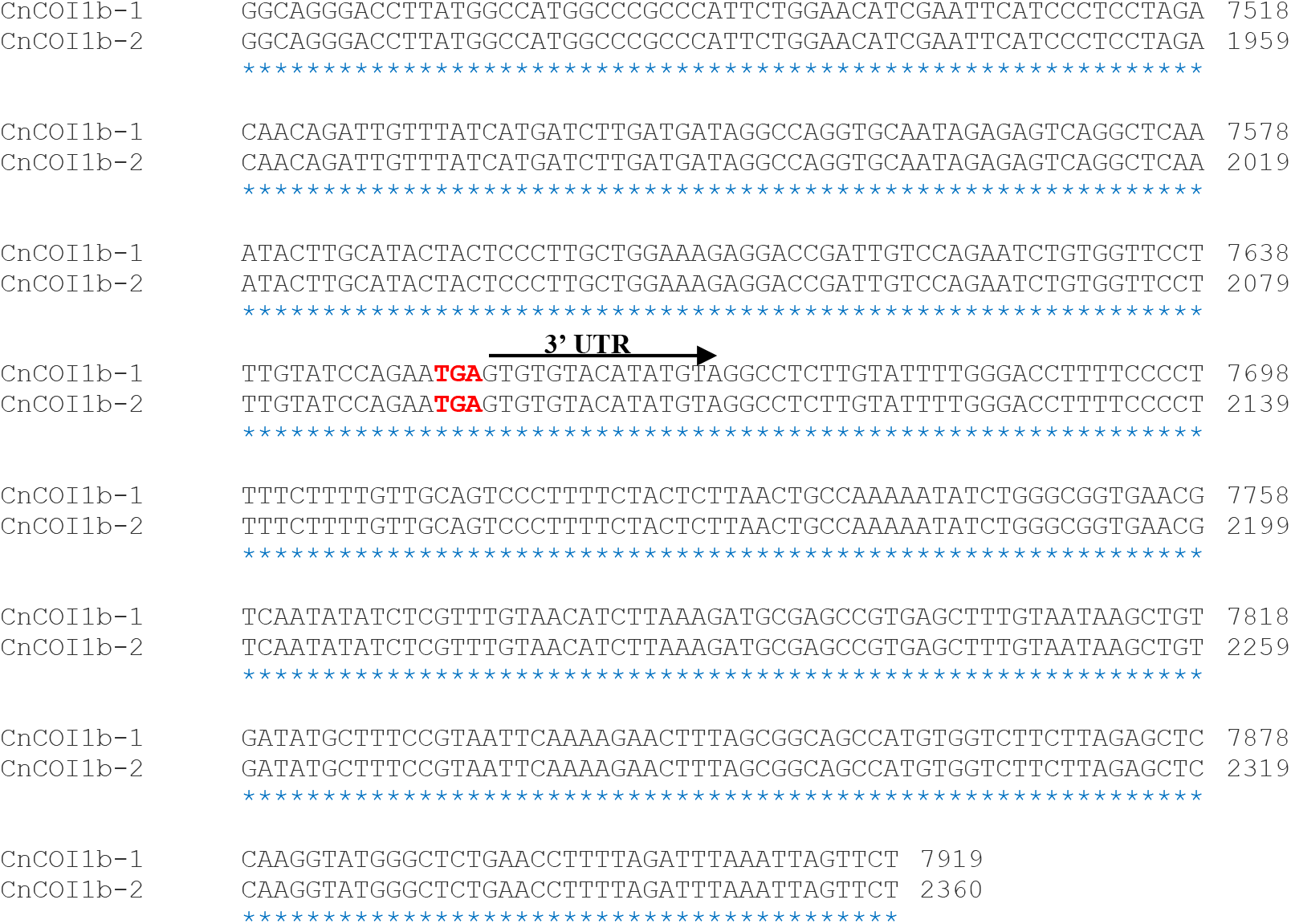
Pairwise nucleotide sequence alignment of CnCOI1b-1 and CnCOI1b-2. The 5’UTR and 3’ÚTR regions are labeled. Start site (green), stop site (red), conserved regions (blue asterisk). The pairwise alignment of 1-5865 bp of CnCOI1b-1 and 1-313 bp of the CnCOI1b-2 were not shown in the figure.

In terms of cDNA sequence characteristic, *E. guineensis* COI1 (XM_010909322.2) showed same number of exons (4) and introns (3), and same length of ORF region (1,743 bp) with CnCOI1b-2. Of all the coconut and representative dicot cDNA sequences of COI1, CnCOI1b-1 had the longest gene length while AtCOI1 had the shortest. In terms of full-length cDNA sequence, CnCOI1b-1 was the longest and the shortest is HbCOI1. Although CnCOI1b-1 had the longest gene and full-length cDNA, it had the shortest ORF. COI from *Aquilaria sinensis* had the longest ORF compared to CnCOI1b, *E. guineensis*, and other dicot species. Furthermore, known COI1 sequences of representative dicot species (**Table 1**) displayed more similarity to CnCOI1b-2 compared to CnCOI1b-1.

### Structural and functional analysis of coconut COI1b protein

The information provided by structural and functional analysis of coconut COI1 can provide the essential and novel details about its biological and cellular function. Though COI1 protein has been extensively characterized in other crops (Xie et al. 1998; Xu et al. 2002; Peng et al. 2009; Liao et al. 2015), its specific function in coconut is yet to be described. The translated amino acid sequences from the two (2) coconut isoforms, CnCOI1b-1 and CnCOI1b-2, span 391 and 580 amino acid residues, respectively (**Table 1**). The molecular weight was predicted to be higher in CnCOI1b-2 (65.5 kDa) as compared to CnCOI1b-1 (44.0 kDa). Both coconut COI1b isoforms are rich in leucine residues (>14%). CnCOI1b-1 protein contains 31.7% helix, 11.8% strand, and 56.5% coiled. On the other hand, CnCOI1b-2 protein contains 35.5% helix, 11.6% strand, and 52.9% coiled. The two isoforms have AMN1 domain and leucine-rich repeat domain (LRR) – 8 LRRs for CnCOI1b-1 and 13 LRRs for CnCOI1b-2 (**Fig 3**).

**Fig 3.**
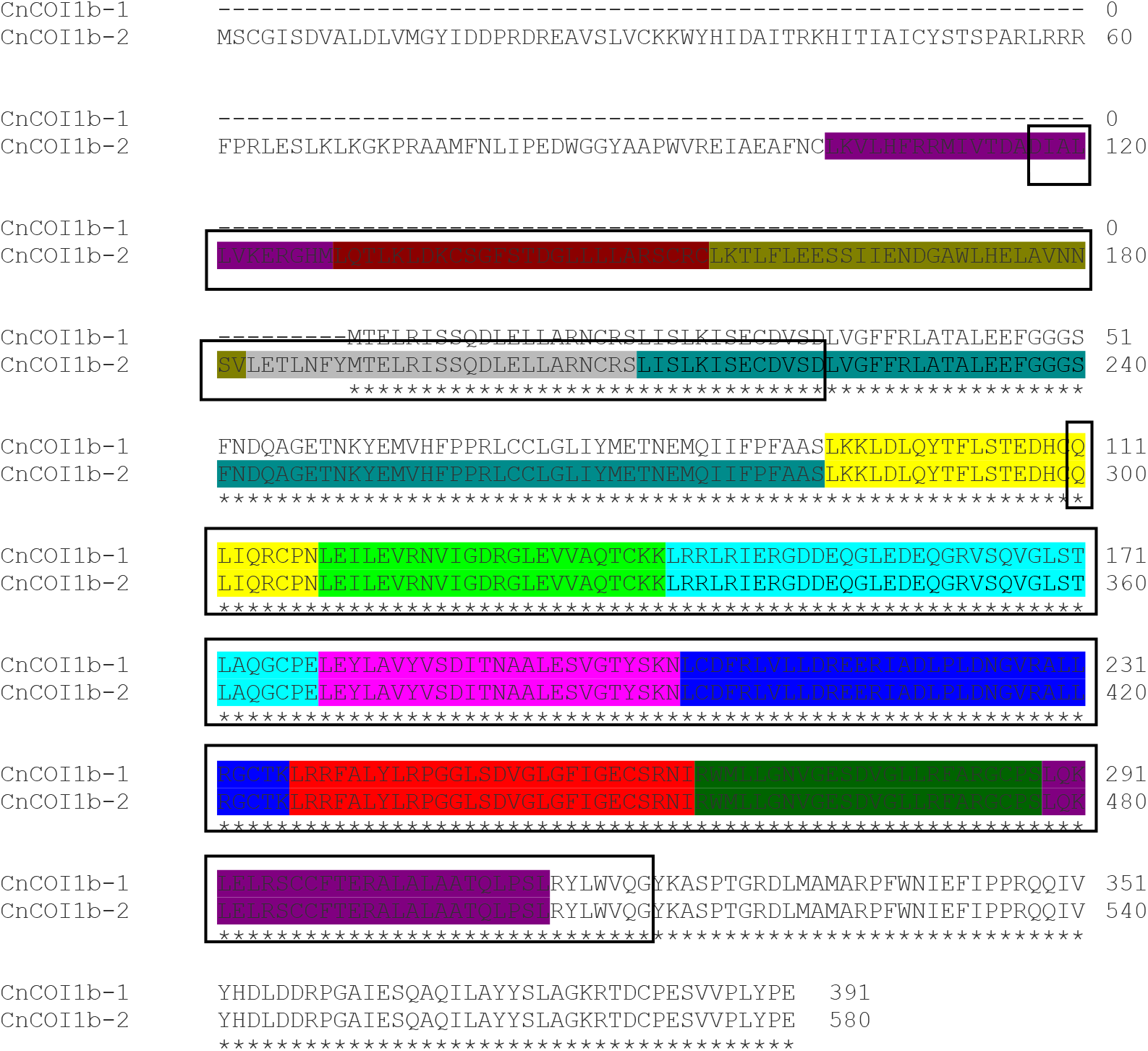
The deduced amino acid sequence of CnCOI1b-1 and CnCOI1b-2. The characteristic motifs are shown as follows: 8 LRRs for CnCOI1b-1 and 13 LRRs for CnCOI1b-2 (highlighted), and AMN1 conserved domain (black box).

Compared to known COI1 protein sequences from other species, CnCOI1b-1 has the lowest molecular weight (44.0 kDa) and the least number of LRR domains (8) (**Table 1**). In terms of secondary protein structure, no significant difference can be observed across COI1 proteins. Only CnCOI1b-1 showed different tertiary structure, having a horse-shoe like shape. All the compared protein sequences showed AMN1 as the top domain hit.

Threading templates used by I-TASSER are based on Z-score or the difference between the raw and average scores in the unit of standard deviation. Based on the result, the highest scoring threading template for both isoforms was the structure of COI1-ASK1 in complex with coronatine and incomplete JAZ1 degron (3ogkB) detected 5X out of 10, from *Arabidopsis thaliana*, classified as protein binding, and expressed in *Spodoptera frugiperda*. The final model for CnCOI1b-1 and CnCOI1b-2 (**Fig 4**) were chosen based on the C-score among the model predictions. The highest structural analog for both isoforms was the Transport Inhibitor Response 1 or TIR1 (2p1oB and 2p1nB), a signaling protein from *A. thaliana* and expressed in *S. frugiperda*.

**Fig 4.**
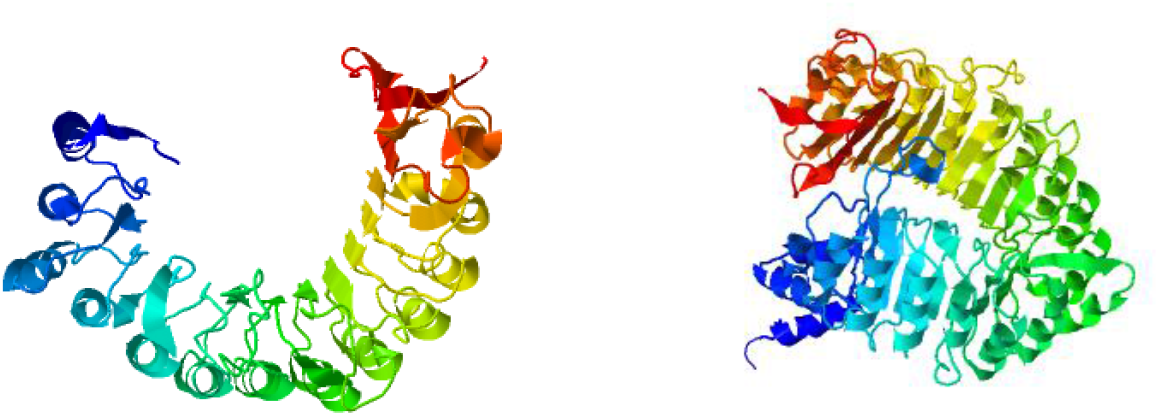
Homology model of CnCOI1b-1 (left) and CnCOI1b-2 (right).

**Fig 5.**
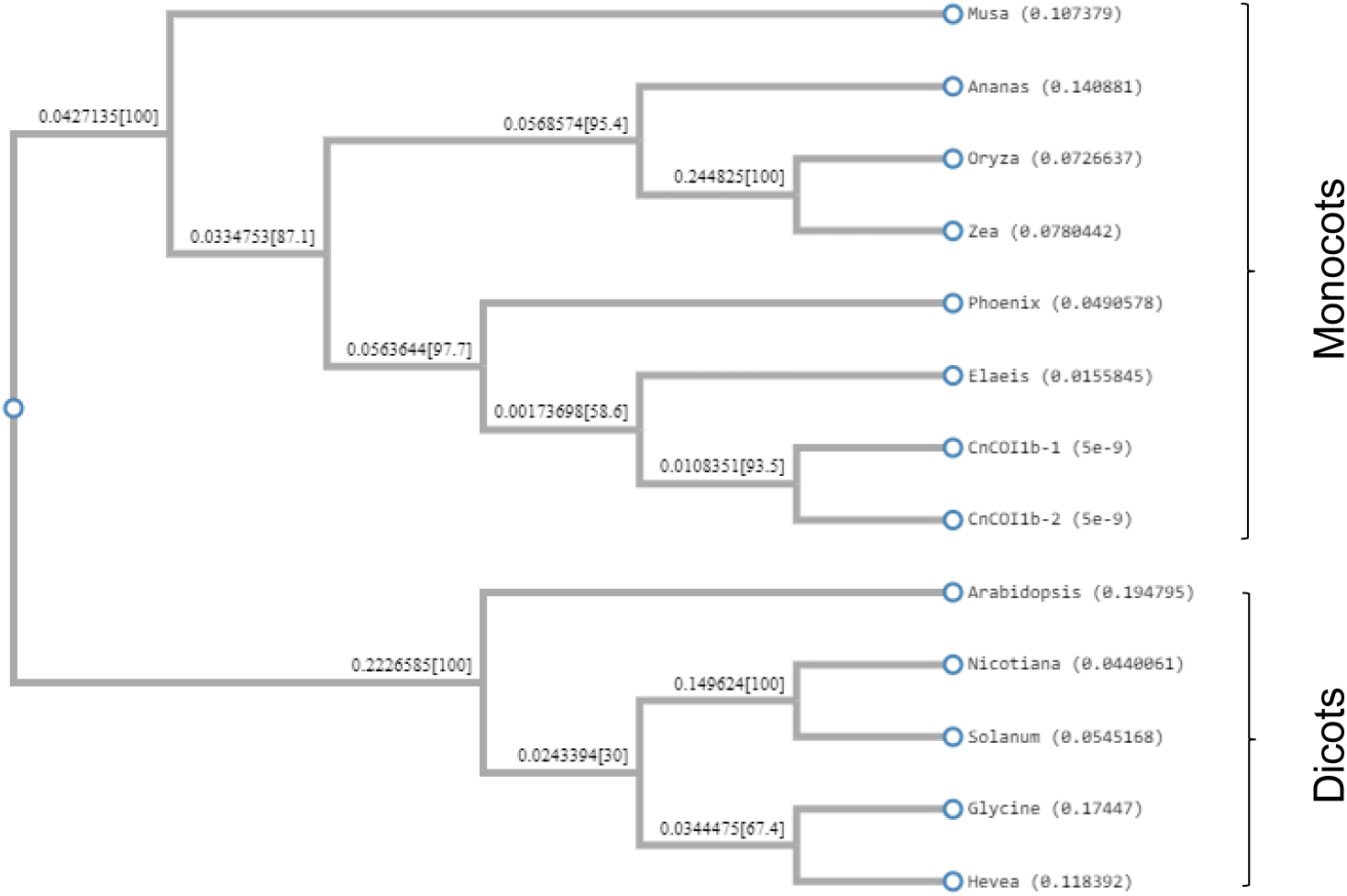
Phylogenetic relationships of the 2 isoforms of COI1b in coconut against known COI1 protein sequences of selected monocot and dicot species. The protein sequences used and their GenBank Accession numbers were: *M. acuminata* (XP_009416210); *A. comosus* (XP_020101802); *O. sativa* (XP_015639870); *Z. mays* (NP_001150429); *P. dactylifera* (XP_008775055); *E. guineensis* (XP_010907624); *A. thaliana* (ABR45948); *N. tabacum* (XP_016479272); *S. lycopersicum* (NP_001234464); *G. max* (NP_001238590); and *H. brasiliensis* (XP_021664949).

All the predicted protein functions for CnCOI1b-1 and CnCOIb-2 were also summarized in **Table 2**. Same ligand, OGK (Coranatine), was detected for both CnCOI1b-1 and CnCOI1b-2. The top enzyme for CnCOI1b-1 was found to be Beta-N-acetylhexosaminidase (EC no. 3.2.1.52), which catalyze the hydrolysis of terminal non-reducing N-acetyl-D-hexosamine residues in N-acetyl-beta-D-hexosaminides, and with active site residue of E123 (Glutamic acid). For CnCOI1b-2, the enzyme membrane alanyl aminopeptidase, which catalyze the removal of N-terminal amino acid from a peptide, amide or arylamide, was found to be present but with no active site residue. The consensus prediction of Gene ontology (GO) terms for CnCOI1b-1 and CnCOI1b-2 were as follows: protein binding, auxin binding, ubiquitin-protein ligase activity, inositol hexakisphosphate binding, ribonuclease inhibitor activity, stomatal movement, jasmonic acid mediated signaling pathway, response to far red light, SCF-dependent proteasomal ubiquitin-dependent protein catabolic process, SCF ubiquitin ligase complex, and Angiogenin-PRI complex. Unique GO terms for each isoform were presented in **Table 2**.

**Table 2.**
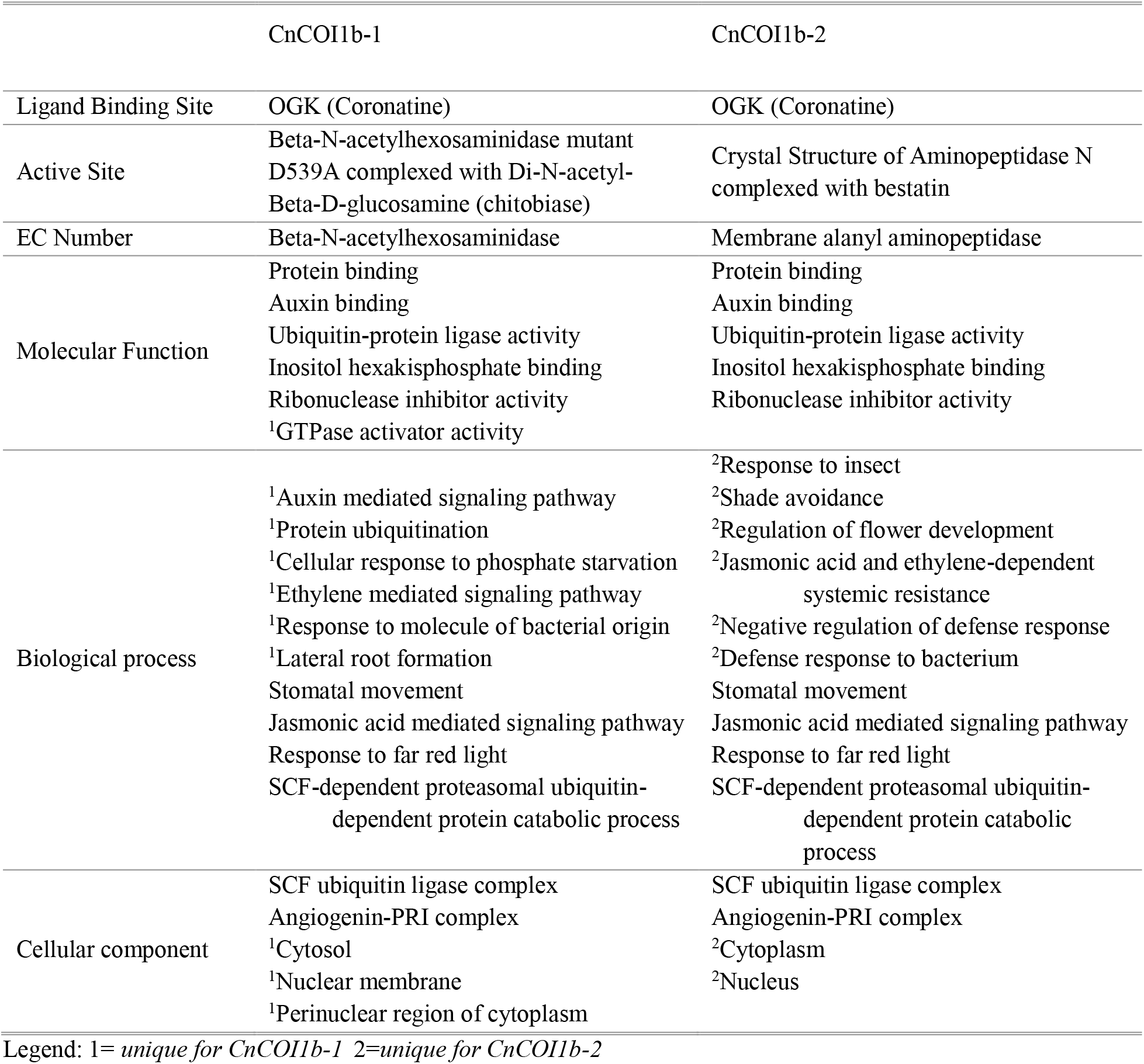
Predicted protein function of CnCOI1 isoforms based using COFACTOR and COACH.

### Homologous alignment and phylogenetic analysis of coconut COI1b

Evolutionary analysis of the coconut COI1b isoforms and their orthologs revealed that these proteins form two (2) subfamilies, one composed of monocots and the other from dicots. The two (2) coconut COI1b isoforms are most related to *E. guineensis* COI1b, which indicate that both have almost similar structure and likely share some same gene function. Two (2) subclades can be observed in the monocot group, one group composed of Arecaceae and the other composed of Musaceae, Bromeliaceae, and Poaceae. COI1 protein in Arecaceae family is more related to Poaceae compared to Musaceae and Bromeliaceae based on the phylogenetic tree.

**Fig 3** Phylogenetic relationships of the 2 isoforms of COI1b in coconut against known COI1 protein sequences of selected monocot and dicot species. The protein sequences used and their GenBank Accession numbers were: *M. acuminata* (XP_009416210); *A. comosus* (XP_020101802); *O. sativa* (XP_015639870); *Z. mays* (NP_001150429); *P. dactylifera* (XP_008775055); *E. guineensis* (XP_010907624); *A. thaliana* (ABR45948); *N. tabacum* (XP_016479272); *S. lycopersicum* (NP_001234464); *G. max* (NP_001238590); and *H. brasiliensis* (XP_021664949).

## Discussion

Previously, *COI1* genes have been cloned and characterized from *Arabidopsis*, *Aquilaria*, rubber, and a few other plant species (Xie et al. 1998; Xu et al. 2002; Peng et al. 2009; Liao et al. 2015), but not from *Cocos nucifera*. The present study is the first report on the cloning and characterization of the gene encoding COI1b protein in coconut. COI1, the first identified F-box protein, is one of the components of the SCF complex that mediates the activation of JA response genes (Xie at al 1998). Mutations in *COI1* in other plant species resulted to JA insensitivity to defense response and physiological processes (Xie et al. 1998; Peng et al. 2009; Liao et al. 2015) which suggest that COI1 is an essential conserved component of JA signaling pathway in plant secondary metabolism.

The multiple sequence alignment of the assembled *COI1b* full-length assembled transcripts against the coconut CATD gene scaffold showed a different orientation for WCT. Possible explanation to this observations are RNA-seq library preparation method (stranded and non-stranded), the treatment (infested or diseased) of the raw read sequence source, and varietal differences. Furthermore, the coconut *COI1b* was found to have two (2) splicing variant – low MW (CnCOI1b-1) and high MW (CnCOI1b-2). Both splicing variants share high similarity with the coronatine-insensitive protein homolog 1b in *E. guineensis*. Long full-length cDNA sequence in *CnCOI1b-1* and short number of amino acid residue suggest that the two mRNAs might be generated by an alternative splicing from a single gene. The occurrence of this kind of post-transcriptional modification has also been reported in human proteosomal modulator subunit, p27 (*PSMD9*) wherein the longer cDNA sequence encodes a shorter polypeptide sequence (Watanabe et al. 1998). Based on transcript data, both splicing variants can be expressed in coconut. Moreover, the differential gene expression response of these coconut COI1b isoforms against insect damage is still subject for further research as previously demonstrated in *Arabidopsis* (Rehrig et al. 2014) and rice (Ye et al. 2012).

Based on striking similarity between COI1b and TIR1, a good structural model (C-score = 0.05, 1.08) for CnCOI1b-1 and CnCOI1b-2 were presented. The observed resemblance between the structures of COI1b and TIR1 provided an indirect evidence to explain the similarity of the signaling pathways for JA and auxin as previously described (Katsir et al. 2008). The conserved LRR domains in CnCOI1b-1 assemble into a solenoid fold which resembles horseshoe shape while CnCOI1b-2 has a spiral structure. The characteristics difference of the LRR domains among the two (2) isoforms might imply that the integrity of the LRR domains is necessary to the structural framework of COI1. Ja-Ile, ligand for the COI-JAZ protein interaction, was reported to bind at the LRR domain of COI1 (Xie et al 1998). The LRR domain of TIR1, highest structural analog for both isoforms, was known to contain inositol hexakisphosphate co-factor and recognizes auxin (Tan et al. 20017).

The identified ligand OGK (Coroatine) of the CnCOI1 was known to have functions associated to plant fertility and defense response, regulation of genes induced by wounding, regulation of jasmonate, among many others (Xie et al 1998, Reymond et al 2000; Feng et al 2003). Some GO terms are unique for each isoform while some are in consensus with one another. Both isoforms almost have the same molecular function except that CnCOI1b-1 has GTPase activator activity which binds to and increases the activity of a GTPase, an enzyme that catalyzes the hydrolysis of GTP. CnCOI1b-1 have biological process associated to auxin mediated signaling pathway while response to insect was only detected in CnCOI1b-2. Both have SCF ubiquitin ligase complex and Angiogenin-PRI complex in their cellular component.

Multiple alignment analysis showed that CnCOI1b had more than 80% sequence identity with COI1 protein of monocots and more than 65% in dicots, which suggest that COI1 is highly conserved specially in monocots. Phylogenetic analysis confirms the high degree of COI1 conservation during the evolution, which reflects the selective pressure imposed by the vital functions of COI1 in plants.

## Conclusion

The *coronatine-insensitive 1* gene (*COI1*) was cloned from coconut using ‘Catigan Green Dwarf’ (CATD) genome sequence assembly as reference in this paper. Two (2) splicing variants were identified - CnCOI1b-1 (low MW) and CnCOI1b-2 (high MW). Although both isoforms revealed different 3D structural models, various physiological and developmental plant processes including defense response to insect and pathogens were conserved. However, differential gene expression response of these coconut COI1b isoforms against insect damage is still a subject of further research. This work may provide the foundation of future research on understanding the role of COI1 in coconut. Results of this study are also expected to assist in the development of new resistant coconut varieties as one of the strategies to address threats in coconut production.

## Acknowledgements

This research was conducted through funds provided by Department of Science and Technology-Philippine Council of Agriculture, Aquatic and Natural Resources Research and Development (DOST-PCAARRD) through the Coconut Project 8 headed by Dr. Hayde F. Galvez. We gratefully acknowledge the Institute of Plant Breeding for allowing us to use the facilities for the conduct of this study. Also, the authors extend their sincerest gratitude to Dr. Susan R. Strickler and Dr. Lukas A. Mueller of the Boyce Thompson Institute for Plant Research (BTI), Ithaca, New York, USA for sharing their expertise in generating the high-quality reference genome assembly. Also, the authors would like to thank Ms. Cynthia R. Gulay for her technical assistance during the conduct of the laboratory experiments.

---

**APPENDIX 1.**
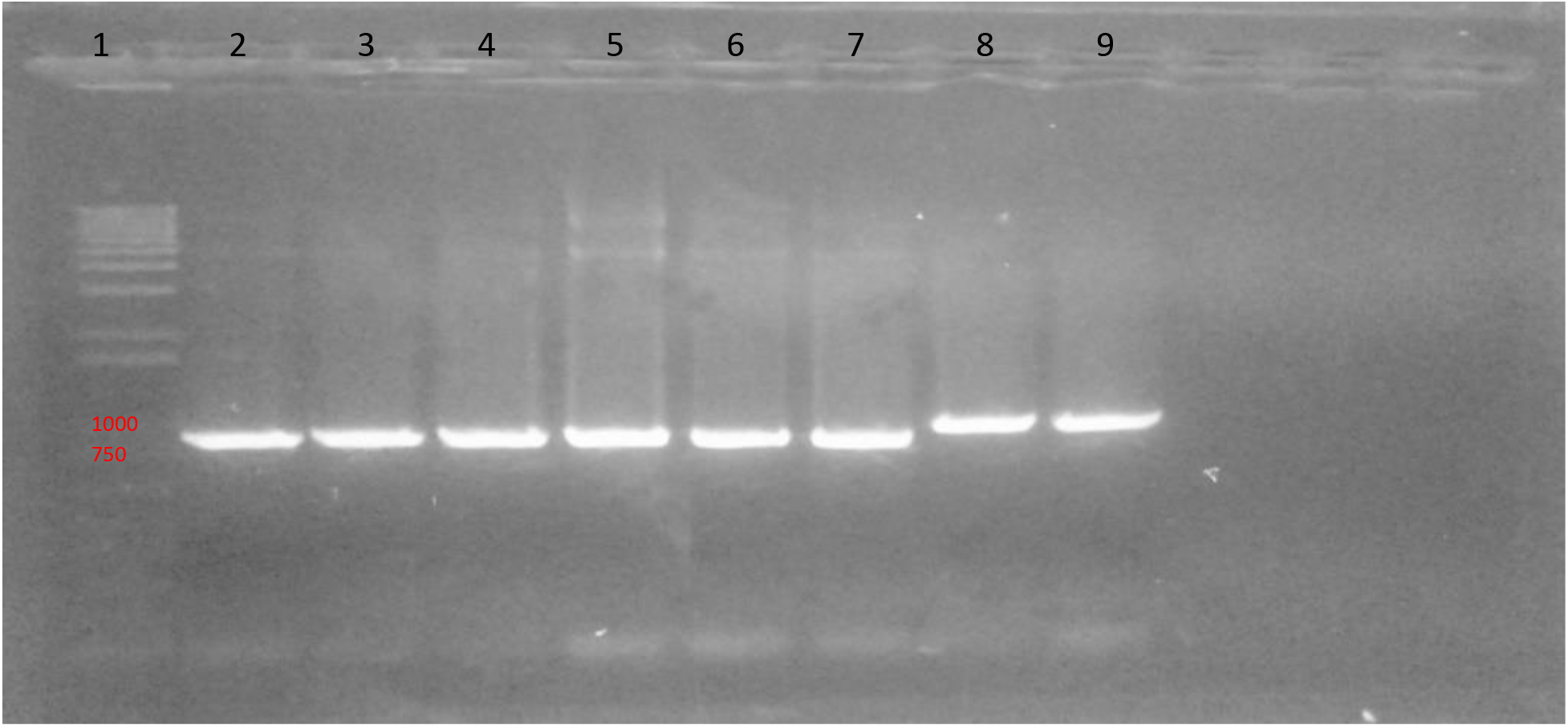
Gel electrophoretogram of insert PCR product: cDNA from healthy CATD leaf samples (Lanes 2-3), CSI-infested CATD leaf samples (lanes 4-7), and genomic DNA of CATD coconut variety (lanes 8-9). A 1kb+ ladder was used.

**APPENDIX 2.**
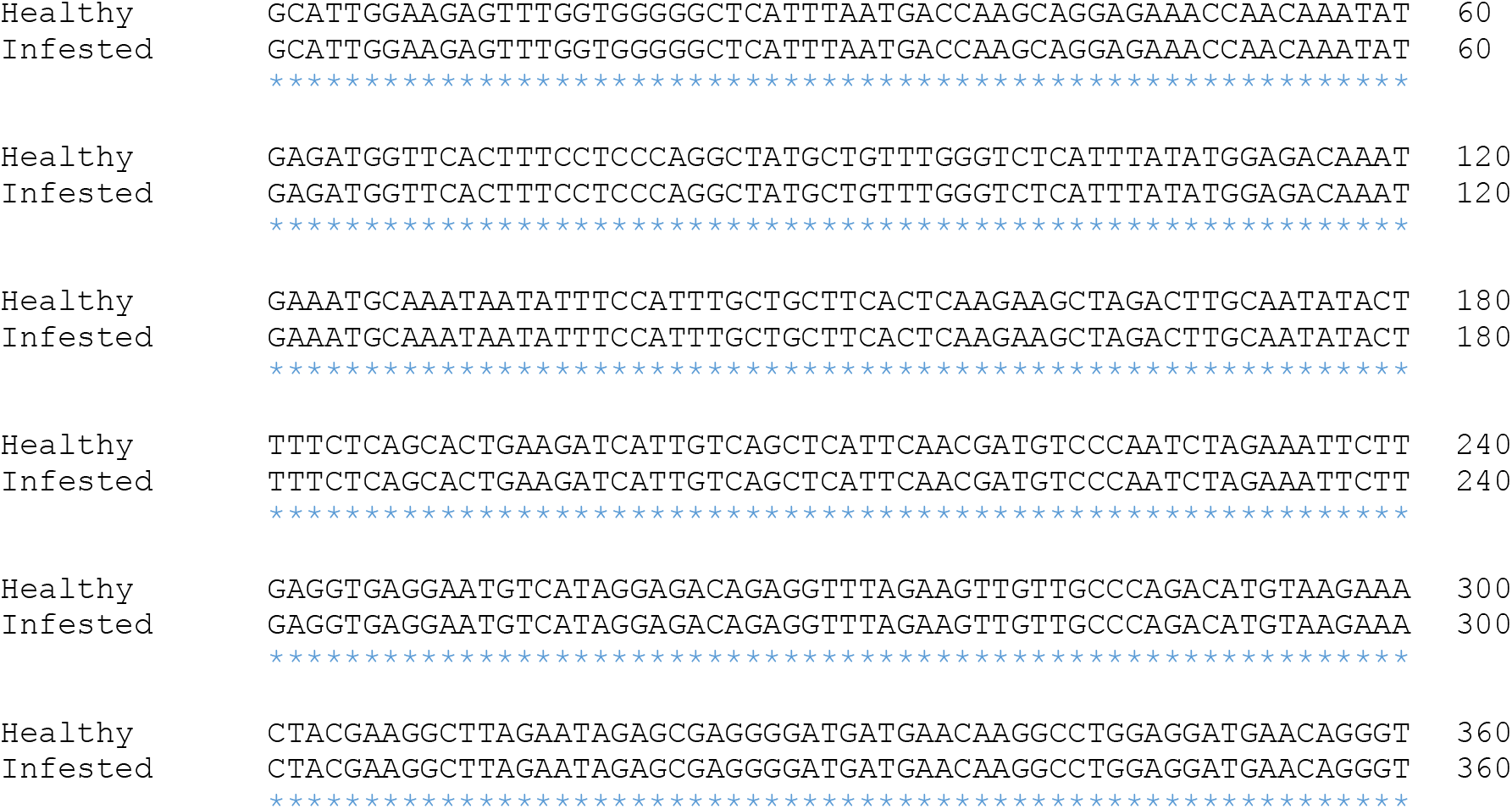
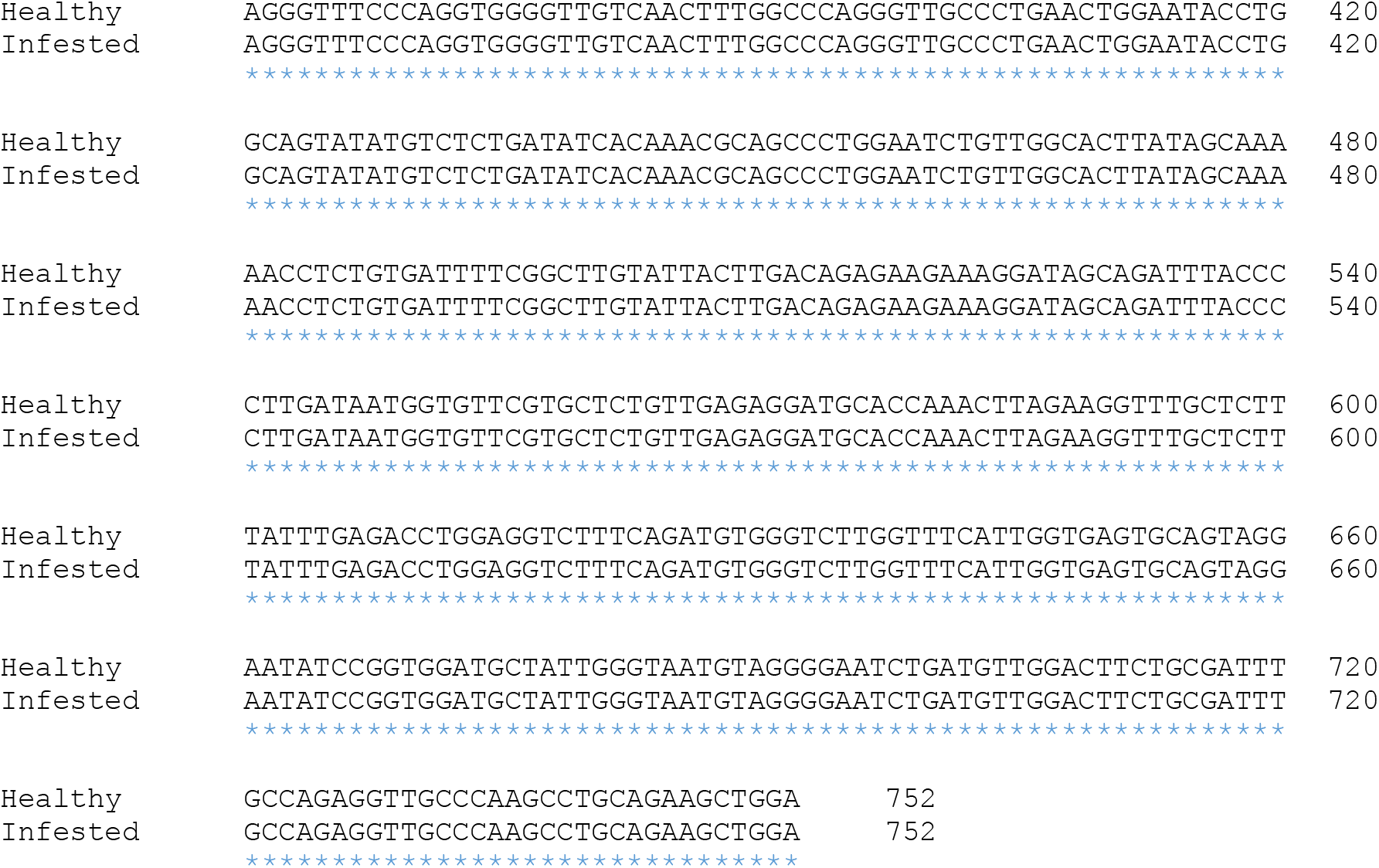
Pairwise alignment of cloned mid-transcript sequences from healthy and CSI-infested CATD leaf samples showed 100% alignment or no sequence variation.

